# Viral tRNA-like structure hijacks host ribosomes for poly(A)-independent translation

**DOI:** 10.1101/2025.10.12.681957

**Authors:** Guoliang Lu, Liming Wan, Yuchen Chen, Ye Li, Yajie Yan, Yan Liu, Jinzhong Lin

## Abstract

Positive-sense RNA viruses often use 3′ tRNA-like structures (TLSs) instead of poly(A) tails to capture the host translation machinery. While TLSs resemble canonical tRNAs and engage host factors, their ability to directly recruit ribosomes has remained unresolved. Here, we present cryo-electron microscopy snapshots of the histidine-accepting TLS (TLS^His^) from tobacco mosaic virus bound to the 60*S* subunit, the 80*S* ribosome, and the 80*S*-tRNA_i_^Met^ initiation complex. Across these states, TLS^His^ is consistently anchored to the L1 stalk of 60*S*, even under cycloheximide treatment, yet remains dynamic on the 40*S* subunit. Structural analysis shows that TLS^His^ transitions from the E-site to an adjacent Z-site to accommodate initiator tRNA. Functional assays show that TLS^His^ preferentially associates with ribosomal subunits over polysomes and can substitute for a poly(A) tail to promote robust cap-dependent translation. Together, these findings reveal how a viral tRNA mimic hijacks host ribosomes to promote poly(A)-independent translation.

## Main Text

In eukaryotic cells, mRNA translation typically requires the 5′ cap and 3′ poly(A) tail to recruit translational factors, such as the eukaryotic initiation factor 4F (eIF4F) complex and the cytoplasmic poly(A)-binding protein (PABPC)^1,2^. The eIF4F complex recognizes the cap structure and recruits the 43*S* pre-initiation complex, forming the 48*S* translation initiation complex that scans the mRNA for the start codon^2–4^. Upon start-codon recognition, the 60*S* ribosomal subunit joins to assemble the 80*S* initiation complex, enabling translation to proceed^3^. The poly(A) tail, present in nearly all human mRNAs, interacts with PABPC to form a closed-loop structure with eIF4F, thereby enhancing translation initiation efficiency^5,6^. In contrast, many viruses have developed mechanisms to bypass cap-poly(A)-dependent translation^7,8^. For example, internal ribosome entry sites (IRESs) in coxsackievirus B3 (CVB3) and hepatitis C virus (HCV) bypass cap-dependent translation, while tRNA-like structures (TLSs) at the 3′ end of viral genomes replace the poly(A) tail^8,9^. Despite their critical roles, the exact mechanisms by which these RNA elements bypass canonical translation remain poorly understood, particularly in the case of TLSs.

TLSs are remarkable RNA elements found at the 3′ ends of many positive-sense plant RNA viruses, such as tymoviruses, tobamoviruses, and bromoviruses^8,10–12^. These structures mimic tRNA functions and can hijack host translation machinery in ways that have long defied understanding since their discovery in the 1970s^13^. TLSs are classified into three major types, TLS^Val^, TLS^His^, and TLS^Tyr^, based on the amino acid they accept^13–15^. Each class adopts a distinct secondary structure that differs from tRNA, demonstrating that tRNA mimicry can occur through diverse architectural strategies^8^. Over the past decades, biochemical studies have shown their aminoacylation activity and affinity for the translation factor eEF1A^13–18^. Structural breakthroughs have revealed how these RNA elements fold into tRNA-like structures and interact with host translation factors. The crystal structure of the TLS^Val^ from turnip yellow mosaic virus (TYMV) has been resolved at atomic resolution, revealing a tightly packed architecture that mimics the canonical tRNA L-shape and explaining its recognition by valyl-tRNA synthetase and eEF1A^19^. More recently, cryo-EM analyses of the TLS^Tyr^ from brome mosaic virus (BMV) uncovered a more flexible, modular structure, likely facilitating binding by tyrosyl-tRNA synthetase and engagement with translation factors^20^. In contrast, the TLS^His^ from tobacco mosaic virus (TMV) has long resisted structural characterization^12^. TLS^His^ lacks the compact tertiary packing observed in TLS^Val^ and TLS^Tyr^, displaying pronounced conformational heterogeneity and a tendency to aggregate^21^. Consequently, TLS^His^ remains the last major TLS class without an atomic-level structure.

Evidence suggests that TLSs exert diverse, virus-specific roles in translation. In TYMV, the TLS^Val^ consists of an upstream pseudoknot domain (UPD) and a following compact tRNA-like domain (TLD). The UPD fine-tunes translation based on local ribosome density, while the TLD enhances translation of genomic RNA in an aminoacylation-dependent manner^22^. In TMV, the TLS^His^ contains multiple pseudoknots (PK1-3) followed by a histidine-accepting TLD, with PK3 proposed to replace the poly(A) tail during cap-dependent translation^23^. In BMV, disruption of TLS^Tyr^ markedly reduces translation of replication proteins, while providing TLS-containing RNA or truncated fragments *in trans* partially restores translation, suggesting a role for TLS in promoting translation initiation or maintaining mRNA stability^24^. The “Trojan Horse” hypothesis proposed that TLS^Val^ could act as a pseudo-initiator tRNA, initiating translation with a valine residue in a cap-independent manner^25^, which provided an important conceptual framework for understanding TLS-mediated translation. However, direct structural evidence for ribosome recruitment by TLSs has remained elusive.

In this study, we resolve this gap by capturing the TLS^His^ from TMV in action on host ribosomes. Using a viral mini-genome mimic, we determined cryo-EM structures of the TMV TLS^His^ bound to the 60*S* subunit, the 80*S* ribosome, and the 80*S*-tRNAi^Met^ initiation complex. These structures reveal that TLS^His^ bridges the 80*S* ribosome by anchoring firmly to the L1 stalk of the 60*S* subunit while dynamically engaging the 40*S* subunit, a binding mode distinct from both canonical tRNAs and the “Trojan Horse” model. Structural analysis shows that TLS^His^ transits from the E-site to the Z-site, a recently defined tRNA-release intermediate, to accommodate P-site initiator tRNA. Functional assays confirm that TLS^His^ can substitute for the poly(A) tail and support robust translation, establishing TLSs as *bona fide* ribosome recruitment elements. Together, these findings provide direct structural and mechanistic evidence that a viral TLS hijacks host ribosomes by exploiting the tRNA-exiting channel to stabilize initiation-competent ribosomes.

## Results

### TLS^His^ engages host ribosomal subunits

The ∼6.4 kb single-stranded TMV genome encodes a replicase and produces two subgenomic RNAs (sgRNAs) for the movement protein (MP) and the coat protein (CP)^26^. All transcripts carry a 5′ cap and a 3′ UTR containing a histidine-accepting tRNA-like structure^27^ (Figure 1A). Whereas the 5′ cap ensures recruitment of canonical initiation factors, how the specialized 3′ architecture contributes to translation remains unresolved.

**Figure 1.**
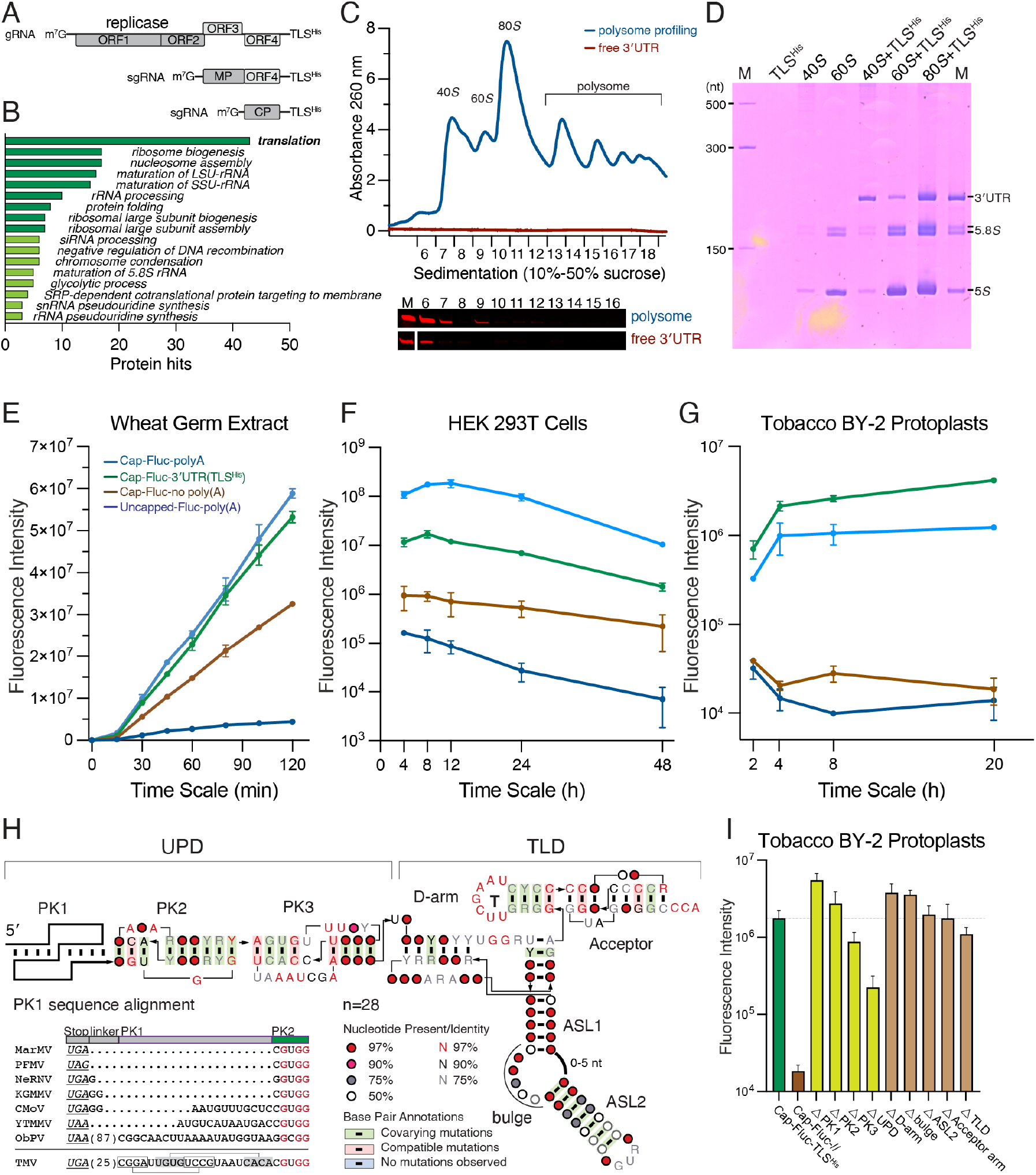
TLS^His^-mediated translation enhancement. (A) Schematic representation of the TMV genome and subgenome, each terminating in a 3′ TLS. (B) RNA pull-down with biotinylated TLS^His^ followed by LC–MS identifies enriched host proteins, predominantly ribosomal components. Polysome profiling (C) and ribosome pelleting (D) reveal preferential TLS^His^ association with free 40*S* and 60*S* subunits, with ribosomal peaks validated by mass spectrometry (Supplementary Data 1). (E–G) Translation efficiency of reporter mRNAs in WGE (E), HEK293T cells (F), and BY-2 protoplasts (G), measured by luciferase activity. (H) Covariation consensus and sequence model of TLS^His^ variants related to the TMV TLS. We surveyed 67 viral strains (37 tobamoviruses and 30 tymoviruses) listed in the latest ICTV release and retained 28 complete 3′ UTR sequences (27 *Tobamovirus*, 1 *Tymovirus*) that contain both the UPD and the TLD for alignment and visualization. The alignment highlights the lack of conservation in PK1 among several viruses. Abbreviations: MarMV, maracuja mosaic virus; PFMV, passion fruit mosaic virus; NeRNV, nemesia ring necrosis virus; KGMMV, Kyuri green mottle mosaic virus; CMoV, cucumber mottle virus; YTMMV, yellow tailflower mild mottle virus; ObPV, Obuda pepper virus; TMV, tobacco mosaic virus. The stop codon is shown in italic and underlined Secondary-structure consensus is annotated underlined. (I) Functional dissection of TLS^His^ shows that both the UPD and TLD contribute to translation enhancement.

To explore whether TLS^His^ directly associates with the translation machinery, we performed a series of biochemical assays, including RNA pull-down coupled with mass spectrometry, polysome profiling, and ribosome pelleting experiments (Figure 1B-D). In RNA pull-downs, a biotin-labeled full-length 3′ UTR immobilized on streptavidin beads was incubated with BY-2 cell lysates, followed by LC–MS analysis. The enriched proteins were predominantly ribosomal components and translation-associated factors, indicating that TLS^His^ engages the host translation apparatus (Figure 1B).

Sucrose gradient polysome profiling with Cy5-labeled TMV 3′ UTR further revealed that TLS^His^ co-sedimented mainly with free 40*S* and 60*S* ribosomal subunits, with little to no signal in 80*S* or polysomal fractions. This suggests that TLS^His^ primarily associates with individual subunits rather than with assembled or elongating ribosome complexes in cell extracts (Figure 1C). In parallel, ribosome pelleting assays using purified components demonstrated that TLS^His^ binds directly and robustly to 40*S*, 60*S*, and also to 80*S* ribosomes *in vitro* (Figure 1D).

Together, these results provide compelling evidence that TLS^His^ physically engages ribosomal subunits across multiple experimental systems, establishing a foundation for functional and structural analyses of how this 3′ RNA element promotes translation.

### TLS^His^ promotes poly(A)-independent translation

To test whether TLS^His^ can substitute for a poly(A) tail, we generated reporter mRNAs containing the TMV 5′ and 3′ UTRs and compared translation with controls, including capped transcripts lacking the 3′ UTR, as well as capped or uncapped transcripts bearing synthetic 100-adenosine poly(A) tails. Translation assays in wheat germ extract (WGE), HEK293T cells, and tobacco BY-2 protoplasts revealed that TMV 3′ UTR-containing mRNAs supported translation as efficiently, or even more so, than polyadenylated mRNAs across all systems tested (Figure 1E-G). Notably, in BY-2 protoplasts, TLS^His^ enhanced translation more than two-fold over the poly(A) tail (Figure 1E), highlighting its adaptation to plant-specific translational machinery. In HEK293T cells, the enhancement was more modest, consistent with previous studies in heterologous mammalian systems ^23,28,29^.

To examine the structural requirements for this enhancement, we created nine mutant constructs with stepwise deletions or disruptions across conserved elements of the 3′ UTR (Figure 1H, I). Translation assays revealed that deletions in the tRNA-like domain (TLD) had minimal impact, while removing PK3 reduced translation by ∼2-fold and removing the entire upstream pseudoknot domain (UPD) caused an ∼8-fold decrease. These results confirm that PK3 is a primary structural determinant of TLS^His^-mediated translation enhancement (Figure 1I). Notably, deletion of PK1 or PK2 led to a slight increase in translation, consistent with previous reports that PK3 in the TMV UPD is critical for replacing the poly(A) tail in cap-dependent translation^23,29^.

Even with PK3 or UPD deletions, expression levels remained significantly higher than those of transcripts lacking the 3′ UTR, suggesting that TLS^His^-mediated enhancement is multifaceted. Both the upstream pseudoknots and the tRNA-like domain work together to enable the TMV 3′ UTR to replace the poly(A) tail and sustain efficient translation initiation.

### Cryo-EM snapshots of TLS^His^ bound to host ribosomes

To address how TLS^His^ engages ribosomes, we attempted to resolve the structure of the ribosome-TLS^His^ complex. Initial complexes assembled with human ribosomes yielded poor TLS^His^ density. We therefore turned to the natural host, isolating ribosomes from *Nicotiana benthamiana* leaves. A synthetic mini-genome RNA containing the capped TMV 5′ UTR, ∼300 nt of coding sequence, and the full 3′ UTR efficiently assembled into ribosome complexes, aided by yeast tRNA to increase heterogeneity (Figure 2A).

**Figure 2.**
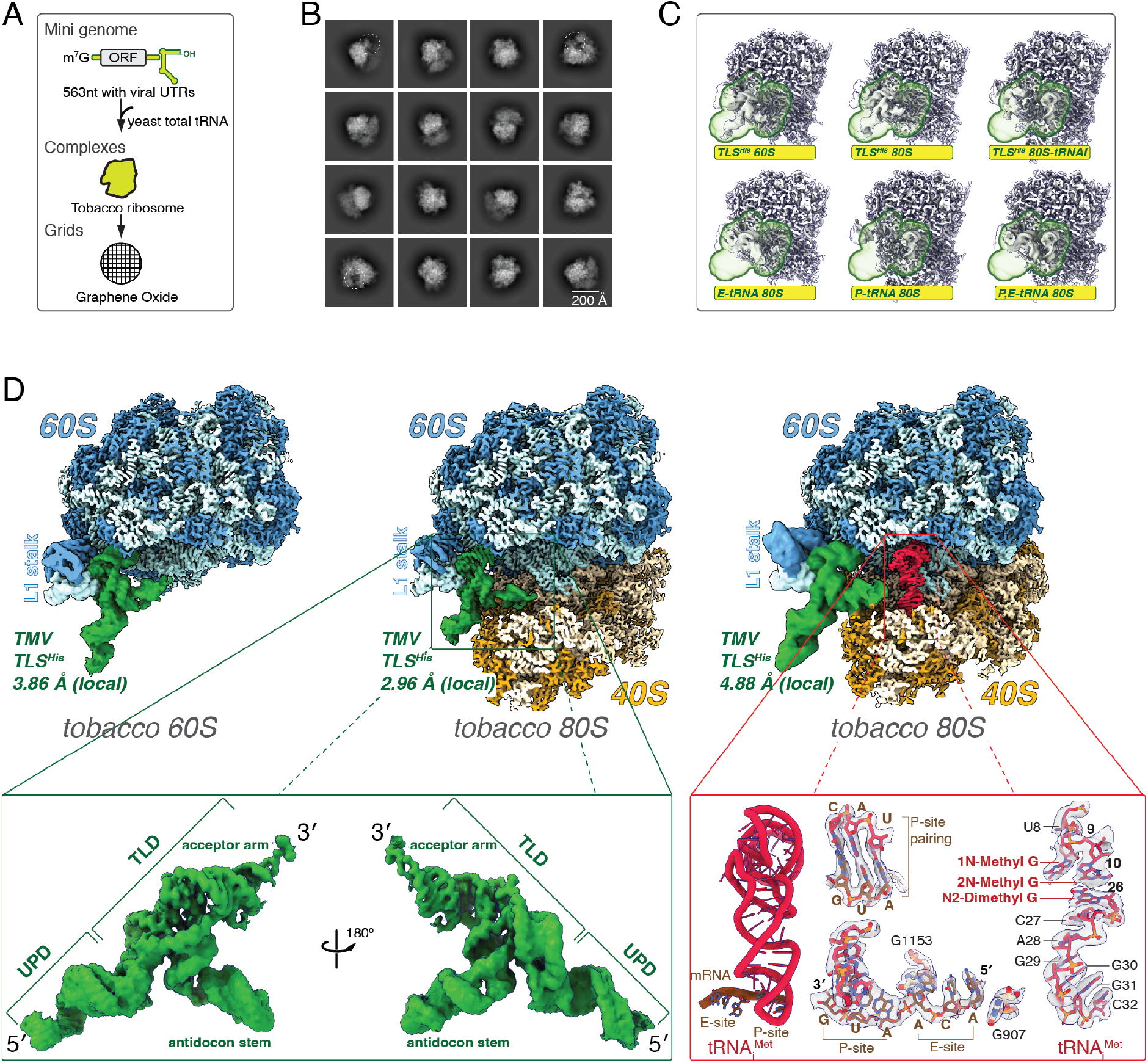
Cryo-EM structures of TLS^His^ bound to host ribosomes. (A) Experimental workflow for TLS^His^-ribosome complex assembly and cryo-EM grid preparation. (B) Representative 2D class averages showing RNA density within tRNA-binding clefts of 60*S* and 80*S* ribosomes. (C) Focused 3D classification around P- and E-sites identifies distinct classes capturing TLS^His^ and P-/E-site tRNA occupancy. (D) Cryo-EM reconstructions of TLS^His^ bound to 60*S* (left), 80*S* (middle), and 80*S* with initiator tRNA (right). TLS^His^ consistently anchors to the L1 stalk. 60*S*/40*S* rRNA and proteins are shown in light/dark blue and yellow/orange; TLS^His^ and initiator tRNA in dark green and red. Inset highlights modular UPD and TLD organization, with mRNA path showing canonical P-site AUG and E-site ACA triplets matching the TMV minigenome sequence.

From 17,264 micrographs, we extracted 3,455,822 particles. Two-dimensional classification revealed strong RNA densities within the tRNA-binding clefts of both 60*S* and 80*S* ribosomes (Figure 2B). Subsequent 3D classification yielded well-defined 60*S* and 80*S* populations. To remove free ribosome signal, particle subtraction combined with 3D classification focused on the E-site and P-site enriched three major states: TLS^His^ bound to the 60*S* subunit, the 80*S* ribosome, and the 80*S* with a P-site tRNA (Figure 2C, Figure S1). Local refinements centered on TLS^His^ and the L1 stalk yielded three distinct structures at 3.86 Å (60*S*-TLS^His^), 2.96 Å (80*S*-TLS^His^), and 4.88 Å (80*S*-tRNA-TLS^His^) (Figure 2D and Figure S1).

In all structures, the TLS^His^ adopts a bipartite T-shaped architecture. The tRNA-like domain folds into a compact L-shape that is buried within the 60*S* E-site, while the upstream pseudoknot domain projects laterally outward from the E-site (Figure 2D). In the 80*S*-tRNA-TLS^His^ complex, the P-site tRNA was unambiguously identified as initiator tRNA_i_^Met^, with clear modification densities (Figure 2D). Within the mRNA channel, the AUG start codon paired with the anticodon of tRNA_i_^Met^, whereas an upstream ACA trinucleotide stacks in the E-site of the 40*S* neck without pairing to TLS^His^. Both features precisely match the TMV 5′ UTR encoded in the synthetic mini-genome (Figure 2D).

### Structural architecture of TLS^His^

Based on the atomic model and structure-guided sequence alignment, TLS^His^ spans 179 nucleotides (residues 26-204) within the 3′ UTR, preceded by a 25-nucleotide linker located immediately downstream of the viral stop codon (UGA). TLS^His^ comprises two major domains, the UPD (residues 26-98) and the TLD (residues 99-204) (Figure 3A). The UPD contains three sequential pseudoknots, PK1 to PK3, while the TLD adopts a tRNA-like fold. Cryo-EM map enabled model building from PK2 to the TLD (residues 48-204), covering ∼87.7% of the TLS^His^. Notably, PK1 is poorly conserved. In several viruses, such as Maracuja mosaic virus (MarMV) and Nemesia ring necrosis virus (NeRNV), the stop codon is immediately followed by a sequence corresponding to PK2 (Figure 1H), suggesting PK1 is dispensable in some lineages.

**Figure 3.**
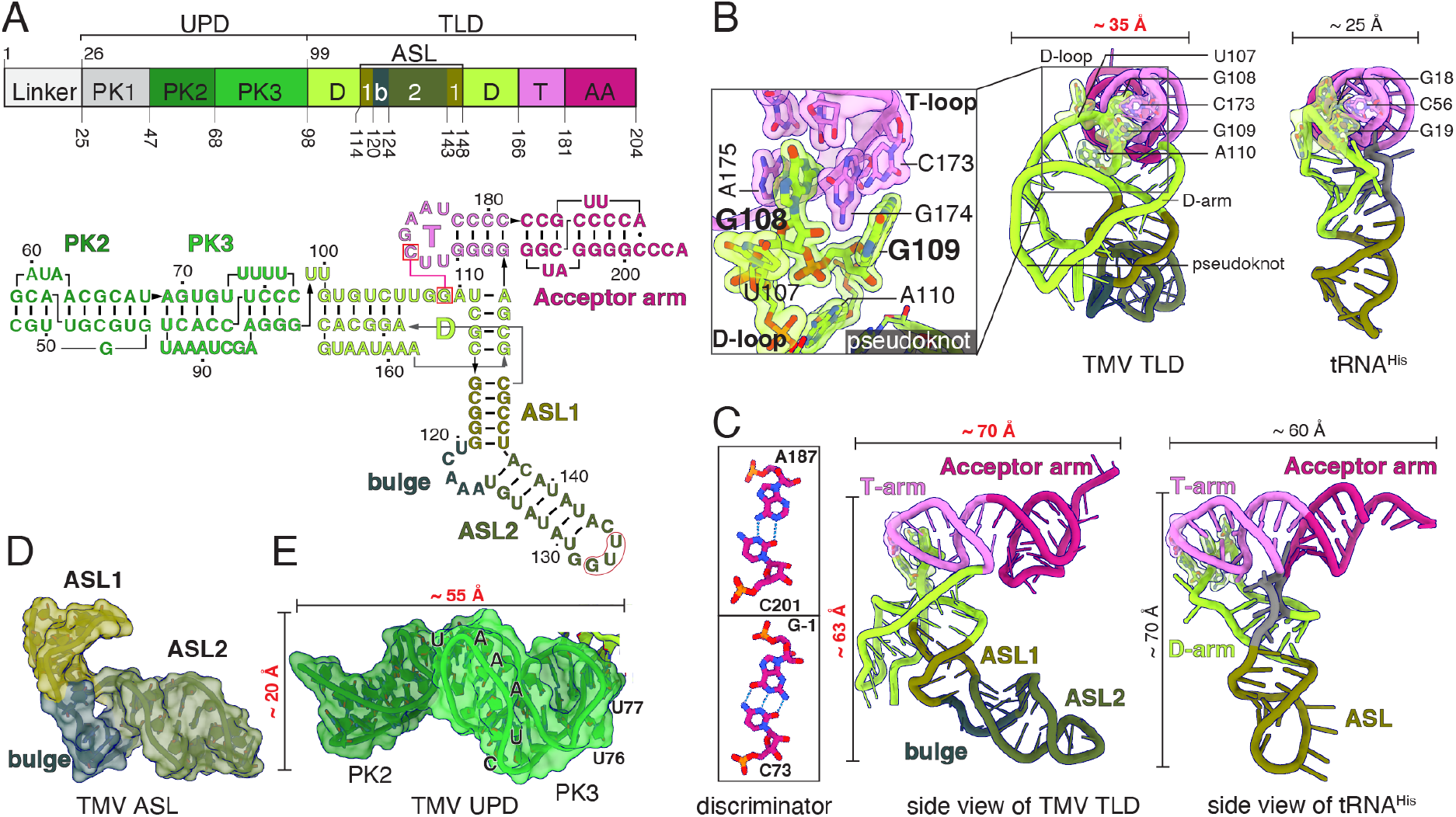
Structural architecture of TLS^His^. (A) Domain organization and secondary structure of TLS^His^. UPD comprises PK1 (disordered, gray), PK2 (dark green), and PK3 (green). TLD contains the acceptor (purple), T (pink), D (light green), and anticodon (olive) arms, with the anticodon arm further subdivided into ASL1, ASL2, and a bulge. (B) Superposition of TLS^His^ and canonical tRNA^His^, highlighting D-arm and elbow rearrangements. D/T-loop interactions mimic the canonical tRNA^His^ fold. (C) Side view of TLS^His^ and tRNA^His^ with the discriminator base pair shown as sticks within the rectangle. TLS^His^ contains an A187:C201 wobble base pair that spatially mimics the canonical G-1:C73 discriminator pair found in tRNA^His^. (D-E) Overview of the ASL (D) and UPD (E) domains. ASL and PKs are shown in cartoons overlaid on the surface.

The TLD forms a L-shaped fold (∼70 Å in length), broadly mimicking the canonical tRNA geometry (Figure 3B, C). Its acceptor arm includes a 3′ pseudoknot resembling TLS^Val^ from TYMV but one base pair shorter (Figure S2). Unlike TYMV, TLS^His^ lacks long-range D-loop interactions, likely reducing structural stability. The discriminator base C201 forms contacts with A187, anchoring the acceptor terminus (Figure 3C). The T-arm closely mirrors canonical tRNA^His^, while the D-arm is replaced by a three-way pseudoknot (Figure 3B). Essential D/T-loop interactions are preserved: G108 (G18 in tRNA^His^) intercalates into the T-loop, and G109 (G19) base pairs with C173 (C56), stabilizing the elbow^30^ (Figure 3B). The anticodon stem is interrupted by a prominent bulge, forming two helices (ASL1 and ASL2) with ASL2 nearly parallel to the acceptor arm (Figure 3C, D). This arrangement positions the D-pseudoknot as a central hub, likely nucleating the overall fold of TLS^His^ fold. Consistent with this role, mutations that disrupt the D/T-loop pseudoknot in related TLS systems severely impair aminoacylation activities^12^.

The flexible UPD limited resolution to PK2 and PK3, which extend ∼55 Å from the D-loop pseudoknot (Figure 3E). These pseudoknots are compactly stacked in sequence, forming a continuous projection outward from the TLD.

### TLS^His^ in action on host ribosomes

TLS^His^ engages host ribosomes by anchoring directly to the L1 stalk of the large subunit (Figure 4A). The ribosomal L1 stalk, a mobile element that guides tRNA movement during translocation, alternates between open and closed states^31,32^ (Figure 4B). TLS^His^ exploits this structural plasticity by coordinating L1 stalk dynamics and repositioning relative to the P-site.

**Figure 4.**
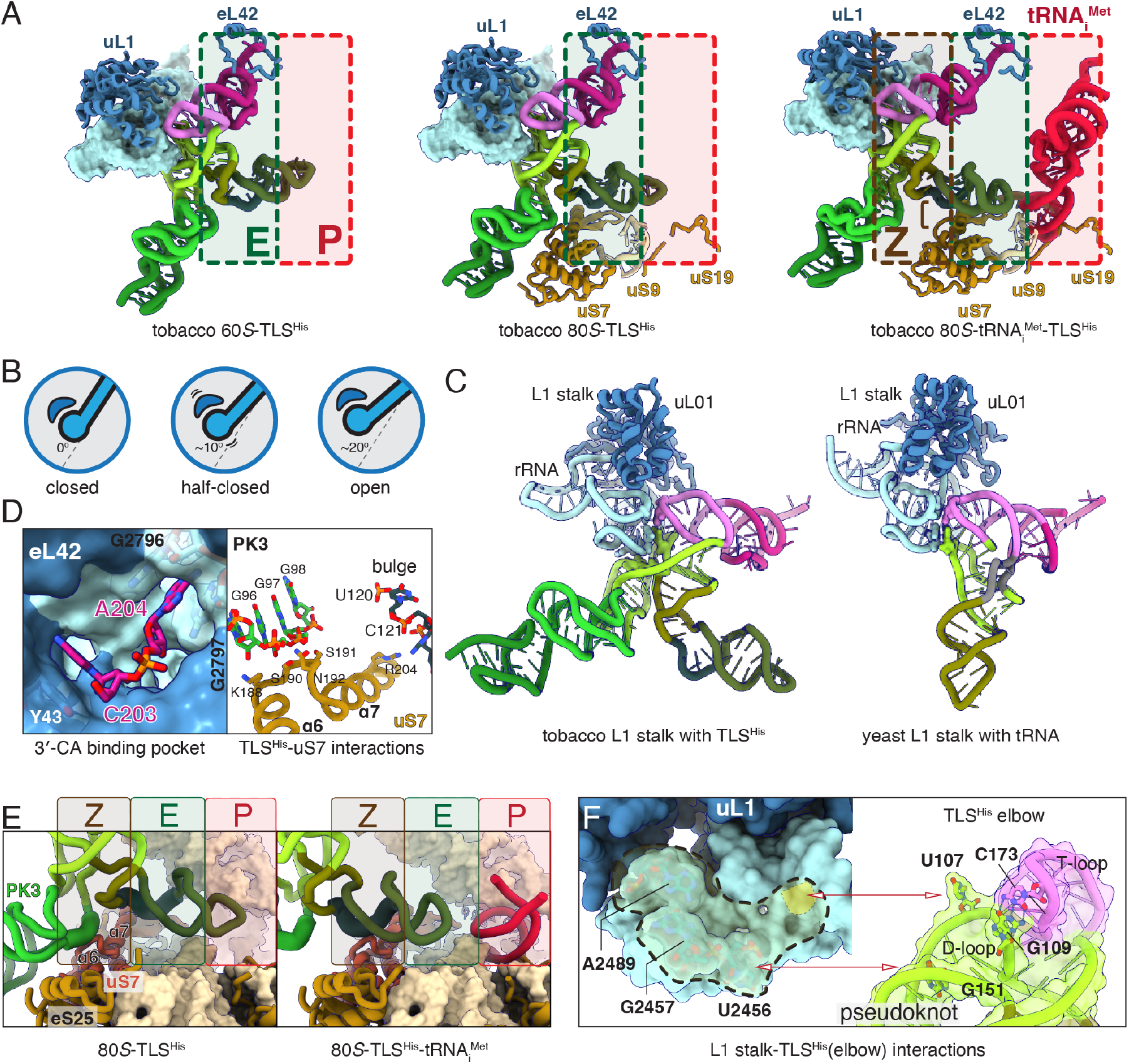
Structural basis of TLS^His^-ribosome engagement. (A) Overview of TLS^His^ complexed with 60*S*, 80*S*, and 80*S*-tRNAi^Met^. TLS^His^ anchors to the L1 stalk across all complexes; In the 60*S*-TLS^His^ and 80*S*-TLS^His^ complexes, ASL2 extends toward the P-site, while shifting outward upon initiator tRNA binding. (B) Cartoon illustration of L1 stalk in closed, half-closed, and open conformations, corresponding to TLS^His^-60*S*, TLS^His^-80*S*, TLS^His^-80*S*-tRNAi^Met^ complexes. (C) TLS^His^ engages 60*S* L1 stalk via D-arm; comparison with canonical tRNA (PDBID: 6gz3). (D) Close-up views of the terminal CCA pocket on the 60*S* subunit and TLS^His^-uS7 contacts on the 40S subunit in the 80*S*-TLS^His^ complex. A204 is sandwiched between 25*S* rRNA bases G2796 and G2797, whereas C203 stacks against the aromatic side chain of eL42-Y43. (E) Comparison of 40S binding patterns between 80*S*-TLS^His^ and 80*S*-TLS^His^-tRNAi^Met^. In the absence of tRNA_i_^Met^, TLS^His^ engages uS7 through both the PK3 and the bulge region of ASL. Upon initiator tRNA binding, TLS^His^ shifts outward, and the interaction is re-established primarily via the backbone of the bulge and the turn between α6-7. (F) Open-book view of the TLS^His^-L1 stalk interface, highlighting three contact networks (dashed circle) with interacting residues shown as sticks.

In the 60*S*-TLS^His^ complex, L1 stalk adopts a closed conformation, with the TLD deeply anchored in the E-site. The terminal A204 inserts into the 28*S* rRNA while the A203 stacks against Tyr43 of eL42 (Figure 4D). The rearranged elbow engages the canonical L1 stalk platform, the G109– C173 base pair stacks against the noncanonical G2457-A2489 pair of L1 rRNA, U107 inserts into the minor groove of the L1 helix, and the sugar of G151 stacks against U2456 (Figure 4C, F). Together, these interactions enlarge the interface to ∼420 Å^2^ compared with ∼280 Å^2^ in canonical tRNAs^32,33^ (Figure 4C).

By contrast, TLS^His^ exhibits conformational flexibility on the 40*S* subunit. Stable binary 40*S*-TLS^His^ complexes could not be captured, but reconstruction of 80*S*-TLS^His^ complex reveals TLS^His^ contacts with two patches on uS7. The loop between α6 and α7 on the 40*S* head engages the PK3 backbone, while the α7 helix at the neck interacts with the bulge loop of the ALS, orienting ASL2 toward the P-site (Figure 4D, E). The TLS^His^ elbow remains anchored to the L1 stalk, and its terminal CCA stays positioned in the E-site, as in the 60*S* complex. Meanwhile, ASL2 protrudes into the P-site, sterically preventing initiator tRNA binding.

In the 80*S*-tRNAi^Met^-TLS^His^ complex, the ribosome adopts a classical nonrotated state with the tRNA in P-site^34^. The L1 stalk shifts to an open conformation, and the bulge of TLS^His^ slides along uS7 to replace PK3 interactions (Figure 4D, E). The bulge-uS7 interaction recapitulates the structural configuration observed for Z-site tRNA stabilized by anticodon-eS25 contacts, described below. TLS^His^ rotates by ∼17° relative to the 60*S* structure and ∼9° relative to the free 80*S* (Figure S3), allowing ASL2 to vacate the P site while stabilizing initiator tRNA. By toggling between closed, half-closed, and open states, TLS^His^ couples L1 stalk dynamics with P-site rearrangements, thereby facilitating the early steps of translation initiation.

### TLS^His^ remains bound despite E-site occupancy

TLS^His^ can likely be aminoacylated with histidine *in vivo*, yet our structures show deacylated TLS^His^ bound to the ribosomal E-site with its acceptor arm, including the CCA terminus, positioned similarly to a classical deacylated tRNA. To test if aminoacylation influences TLS^His^ binding, we instead employed the antibiotic cycloheximide (CHX) to sterically block the E-site, taking advantage of its well-characterized ability to prevent deacylated tRNA from occupying the E-site of ribosomes^35^.

Polysome profiling performed in the presence of CHX revealed that TLS^His^ continued to co-sediment with 40*S* and 60*S* fractions, indicating that its association with large ribosomal subunits does not require classical insertion of the CCA end into an unoccupied E-site (Figure 5A). Ribosome pelleting assays further confirmed that CHX had no detectable impact on TLS^His^ binding to the 60*S* subunit, nor did treatment with puromycin (Puro) or homoharringtonine (HHT), which targe the 60*S* A-site^36^ (Figure 5B).

**Figure 5.**
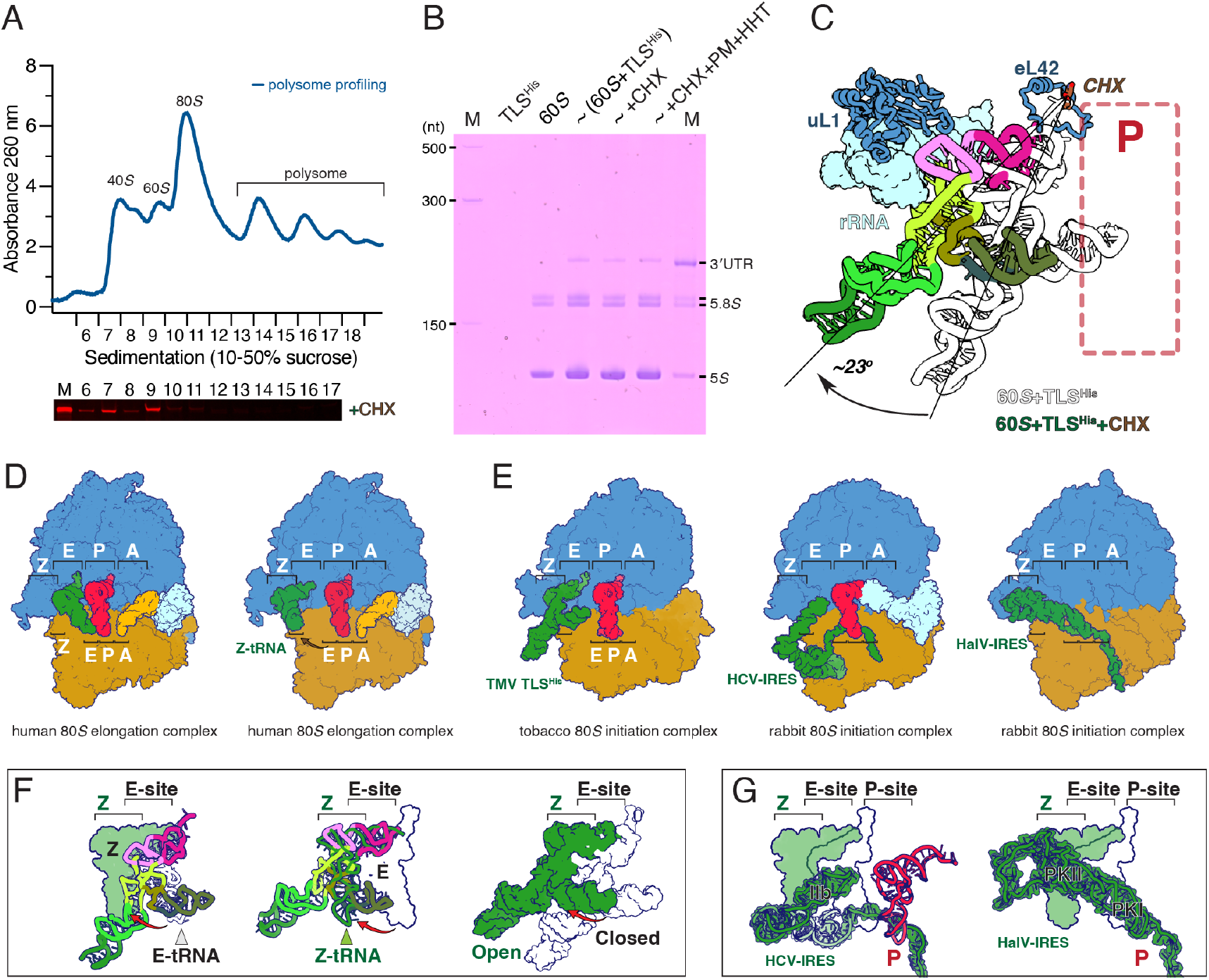
TLS^His^ remains bound to 60*S* in the presence of CHX. (A) Polysome profiling shows CHX does not disrupt TLS^His^ association with free 60*S* subunits. (B) Ribosome pelleting confirms stable TLS^His^-60*S* binding despite E-site occupation. (C) Comparison of CHX-bound and CHX-free TLS^His^ aligned on 25*S* rRNA (L1 stalk, residues 2454-2500) reveals an outward rotation of ∼23°. (D) Cryo-ET structures capture deacylated tRNA in the intermediate Z-site of the human 80*S* ribosome (PDBID: 9azs-E-tRNA; 9b0o-Z-tRNA). (E) Structural comparison of TMV TLS^His^ with HCV and HalV IRESs highlights a shared strategy, the ribosomal tRNA exiting channel is broadly repurposed as a regulatory hub where viral RNAs stabilize initiation-competent ribosomal states (PDBID: 4ujc-HCV-IRES; 7a01HalV-IRES). Viral RNA elements or E/Z-site tRNA are shown in dark green, P-site tRNA in red, A-site tRNA in orange, and eEF1A/eIF5B in cyan. (F) Comparison of E-site and Z-site tRNAs with TLS^His^. The closed TLS^His^-60*S* complex resembles an E-site tRNA, whereas the open conformation captured with 60*S*--CHX and 80*S*-tRNA_i_^Met^ mimics a Z-site tRNA. (G) Comparison of viral IRESs with E/Z-site tRNAs. The HCV type III IRES engages the E-site via domain IIb, while the HalV IRES spans from the Z-site to the P-site. P-, E-, and Z-site tRNAs are shown in red, white, and green, and IRES domains in green cartoon overlaid on the surface.

To explore this interaction structurally, we determined the cryo-EM structure of the TLS^His^ with 60*S* under CHX treatment (Figure S5). The resulting map clearly resolved CHX occupying the E-site, engaging conserved 25*S* rRNA nucleotides and overlapping the position normally occupied by the A76 of a deacylated tRNA, consistent with its established mechanism of E-site blockade (Figure S4A)^35^. Remarkably, despite this blockade, TLS^His^ remained stably anchored to the L1 stalk via its elbow region. This partial release of anchoring allowed the L1 stalk-TLS^His^ module to swing outward by ∼23° relative to the CHX-free 60*S* complex, and ∼6° relative to the 80*S*-TLS^His^-tRNA_i_^Met^ complex (Figure 5C, S4C). This outward repositioning shifts ASL2 away from the P-site, eliminating potential steric clashes with initiator tRNA during subunit joining. Superimposition of the L1 stalk revealed that its interface with the TLS^His^ elbow was essentially unchanged (Figure S4B). Only the terminal CA dinucleotide of TLS^His^ became displaced and unresolved, reflecting loss of stable CCA stacking.

Comparison with ribosome-bound Z-site tRNA^37–41^, recently described as release intermediates, reveals a striking parallel. Translation inhibition by CHX leads to robust accumulation of tRNA at the Z-site, closely resembling our observations^41^. Thus, TLS^His^ exploits a similar site, but instead of marking release, it stabilizes a configuration that primes translation initiation (Figure 5D, F). This strategy also mirrors other viral RNAs that utilize the E-site to promote initiation. Type III and IV IRESs, such as those in HCV^42^, halastavi árva virus (HalV)^43^, and cricket paralysis virus (CrPV)^44^, extend structural domains into the E-site to either position the initiator tRNA or bypass it entirely via pseudoknot-mediated decoding, often in coordination with the L1 stalk (Figure 5E, G). These viral RNAs repurpose the E-site to facilitate initiation, leveraging structural mimicry and ribosomal anchoring to control translation.

Together, these findings demonstrate that TLS^His^ binding does not rely on canonical CCA insertion. Instead, steric blockade of the E-site, whether by aminoacylation or CHX, shifts E-site TLS^His^ into a Z-tRNA-like configuration that favors 60*S* joining and 80*S* assembly. Converging strategies across TLSs and viral IRESs suggest that the ribosomal E-site, traditionally viewed as an exit pathway for deacylated tRNAs, has been broadly repurposed as a regulatory hub for viral RNAs to stabilize initiation-competent ribosomal states.

### A model for TLS^His^-driven poly(A)-independent translation

Our combined biochemical and structural data support a model in which TLS^His^ enables TMV RNAs to achieve robust translation without a poly(A) tail by directly recruiting and positioning ribosomal subunits (Figure 6).

**Figure 6.**
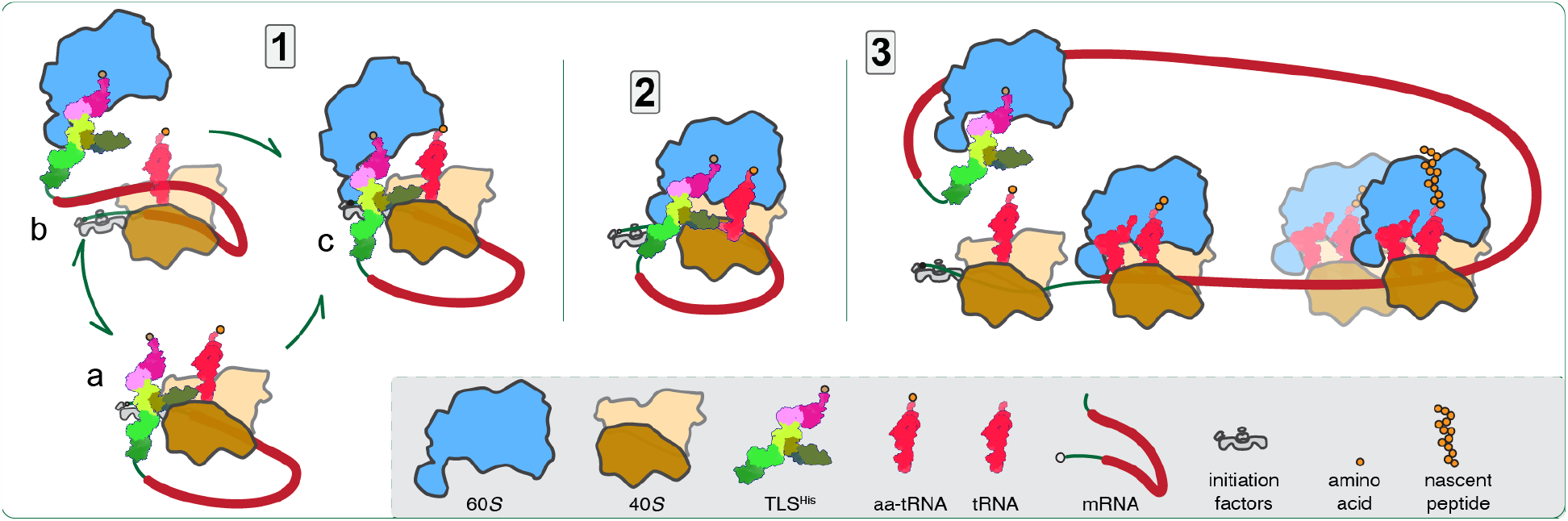
Model of TLS^His^-mediated translation enhancement. Schematic illustrating TLS^His^ (color-coded as in Figure 3) promoting cap-dependent translation without a poly(A) tail. TLS^His^ can associate with both ribosomal subunits. When bound to the 40*S* subunit (a), TLS^His^ facilitates recruitment of the 60*S* subunit (c), enabling assembly of an initiation-competent 80*S* ribosome. Alternatively, TLS^His^ can first engage the 60*S* E-site (b), where its TLD anchors to the E-site, thereby pre-positioning the large subunit for joining with the 48*S* pre-initiation complex to form the 80*S* initiation complex. (2) Once the 80*S* initiation complex is assembled, TLS^His^ is displaced upon A-site tRNA accommodation and nascent peptide elongation. (3) The released TLS^His^ is then available to capture new 40*S* or 60*S* subunits, supporting successive rounds of initiation.

The TMV genome carries a 5′ cap and thus initiates translation via the canonical cap-dependent pathway. Functional assays demonstrate that TLS^His^ can effectively substitute for a poly(A) tail in promoting cap-dependent translation, with the UPD, particularly PK3, playing a role analogous to the poly(A)-PABP-eIF4F system that circularizes host mRNAs and stabilizes initiation complexes. In place of this typical host strategy, TMV employs TLS^His^ as an RNA-encoded mechanism to drive translation.

Our cryo-EM structures provide a mechanistic basis for this substitution. TLS^His^ stably anchors to the 60S subunit via its TLD, inserting into the E-site and extensively engaging the L1 stalk. This association likely enables TMV RNAs to sequester free 60*S* subunits, ensuring a local pool of large subunits that are readily available for rapid initiation on viral transcripts.

Concurrently, TLS^His^ positions PK3 to contact the 40*S* head, particularly via phosphate backbone interactions with uS7, effectively bridging the large and small subunits. This arrangement pre-positions a 60*S* subunit adjacent to the scanning 48*S* complex on the TMV 5′ UTR, lowering the energetic barrier for subunit joining and accelerating assembly of the 80*S* initiation complex.

This model is further supported by our polysome profiling data, which show enrichment of TLS^His^ with free 40*S* and 60*S* subunits but little association with 80*S* or heavy polysome fractions. Such a distribution suggests that TLS^His^ binding to 80*S* ribosomes is transient, consistent with a scenario in which TLS^His^ promotes subunit joining but disengages once elongation commences. After each round of initiation, TLS^His^ would then be free to bind another 60*S* and/or 40*S* subunit, perpetuating the cycle and driving successive rounds of translation initiation on the same viral RNA.

The mobility of TLS^His^ on the ribosome, anchored through interactions with the L1 stalk on the 60*S* and phosphate backbone contacts with the 40*S*, likely facilitates this process. By loosely bridging ribosomal subunits, TLS^His^ remains dynamic enough to accommodate incoming initiator tRNAs. During elongation, displacement from the Z-site may occur as the deacylated P-site tRNA translocates into the E-site, freeing TLS^His^ to capture additional 60*S* subunits for new initiation events.

Through this mechanism, TLS^His^ not only replaces the classical poly(A)-mediated recruitment machinery but also actively organizes ribosomal subunits, endowing the TMV TLS element with the capacity to enhance translation *in cis*.

## Discussion

Our study provides the first high-resolution structural characterization of a viral TLS bound to host ribosomes. This not only yields the first ribosome-engaged structure of TLS^His^, but also completes the structural landscape of the three major viral TLS classes: TLS^Val19^, TLS^Tyr20,45^, and TLS^His^, each now defined in molecular detail (Figure S5). Beyond its previously known interaction with aminoacyl-tRNA synthetases, TLS^His^ is here visualized directly in complex with ribosomal subunits, establishing a new paradigm for TLS function. Together with our biochemical assays, these findings illuminate how TLS elements can act directly in translation, extending their established roles in replication and genome packaging.

TLS^His^ exhibits a divergent adaptation of the canonical tRNA fold. Although it retains the overall L-shaped geometry, it incorporates distinctive local features absent from both canonical tRNAs and other viral TLSs. The most notable is the replacement of the D-arm with a three-way pseudoknot junction^46^, in which a conserved GG dinucleotide engages the T-loop to mimic the canonical D/T-loop pairing of authentic tRNA^His12^. Moreover, the anticodon arm is disrupted by a bulge that distorts the anticodon stem loop, raising questions about how these deviations affect recognition by HisRS. Unlike canonical tRNA^His^, which typically requires a G-1:C73 base pair and a free monophosphate at G-1 for efficient recognition^30^, TLS^His^ exhibits an A187:C201 pairing. Earlier biochemical data showed that mutation of G-1 to A-1 does not abolish aminoacylation^30^. These findings suggest that HisRS recognition is more permissive than previously appreciated, and future structural studies of the TLS^His^-HisRS complex will be essential to understand how such noncanonical features are accommodated.

Our data reveal that TLS^His^ replaces the classical poly(A)-PABPC-eIF4F circuit through a modular RNA mechanism. Functional assays showed that the UPD, particularly PK3, plays a role analogous to a poly(A) tail in bridging events that promote translation. This conclusion aligns with previous deletion analyses that established PK3 as critical for poly(A)-independent enhancement^23,29^. Our structures now provide a mechanistic basis by showing PK3 contacts with uS7 on the 40*S* head, positioning TLS^His^ for productive engagement with the P-site. This modular architecture recalls SERBP1, recently implicated in bridging ribosomal subunits *in vivo*, and underscores the evolutionary flexibility of 3′ RNA modules^41^. Strikingly, related tobamoviruses such as hibiscus latent Singapore virus achieve similar outcomes using internal poly(A) tracts instead of structured UPDs^47^, illustrating parallel evolutionary solutions for maintaining translation without a canonical poly(A) tail.

Across all captured complexes, TLS^His^-60*S*, TLS^His^-80*S*, TLS^His^-80*S*-tRNA_i_^Met^, and 60*S*-TLS^His^ under CHX treatment, the TLS consistently anchors within the E-site while sampling closed, half-closed, and open stalk conformations. This dynamic engagement mirrors the natural motions of the L1 stalk during tRNA translocation, but here TLS^His^ exploits them to facilitate initiation. The TLS^His^ elbow establishes an unusually extensive interface with the L1 stalk^32,36^, and this interaction persists even when the CCA end is destabilized, suggesting that aminoacylation is dispensable for ribosome binding. Strikingly, this conformation parallels the recently described “Z-site” tRNAs^38– 41^. The release intermediates positioned adjacent to the E-site and terminal A76 no longer stacks within 28S rRNA^41^. Both the displaced CCA terminus and the relative orientation resemble TLS^His^, suggesting that the site normally sampled by exiting tRNAs is exploited by viral TLSs to stabilize a configuration that primes initiation.

Several viral RNAs similarly exploit the ribosomal E-site. The HCV IRES inserts its IIb and IIf domains into the E-site to position initiator tRNA^42^, while the HalV IRES binds the 80*S* directly and uses a downstream pseudoknot to bypass canonical initiation^43^. Other examples include cap-independent translation enhancers (CITEs), such as the turnip crinkle virus 3′ CITE, which interacts with the 60S subunit to stimulate initiation^48^. TLS^His^ thus joins a broader repertoire of structured viral RNAs that use the E-site as a regulatory hub.

From an evolutionary perspective, direct ribosomal recruitment by TLS^His^ likely provides TMV with a translational advantage. By pre-binding free 60*S* subunits and bridging them to the 5′ engaged 40*S* complex, TLS^His^ may accelerate 80*S* assembly and thereby favor the efficient translation of viral proteins. Moreover, association with 60*S* could bias ribosome availability toward viral RNAs, particularly under cellular stress or host translation shutoff. Such strategies exemplify a general evolutionary principle: structured RNA elements evolve to co-opt host translation machinery, ensuring robust viral protein synthesis even in adverse environments.

Collectively, our findings establish TLS^His^ as a sophisticated RNA-based alternative to poly(A)-mediated translation enhancement, illustrating how viruses harness tRNA mimicry not only for replication and encapsulation but also to directly remodel host translation. Whether this strategy of direct 60*S* recruitment extends to other TLS classes, such as TLS^Val^ or TLS^Tyr^, remains an open question deserving further study. More broadly, these insights highlight opportunities to engineer synthetic mRNAs that incorporate TLS modules to achieve stable, high-efficiency translation, a concept with potentially transformative applications in mRNA vaccine and therapeutic design.

## Acknowledgements

We thank Dr. Jinbiao Ma for valuable discussions, and Li Li for technical assistance. We are grateful to the Cryo-Electron Microscopy Core Facility of the School of Life Sciences, Fudan University, and to Dejian Zhou for support with cryo-EM data collection. We also thank Chuanwei Yang for guidance on *Nicotiana benthamiana* cultivation, Guohui Xie and Yijing Zhang for providing the tobacco BY-2 cell line, and Lin Huang for assistance with LC-MS analyses.

This work was supported by the National Key Research and Development Program of China (Grants 2023YFC2604303 and 2023YFC2306404 to JL), the National Natural Science Foundation of China (Grants 32130063, 32471349, and 32371352 to JL), the Shanghai Municipal Science and Technology Commission (Grant 24HC2810600 to JL), and the Innovation Program of the Shanghai Municipal Education Commission (Grant 2021-01-07-00-07-E00074 to JL) and the Zhangjiang mRNA Innovation and Translation Center. We sincerely acknowledge all sources of funding.

## Author contributions

J.L. conceived and supervised the project. Y.C., L.W., and G.L. prepared RNA constructs. G.L., Y.C., and Y.Li. purified tobacco ribosomes. G.L. performed ribosome pelleting, polysome profiling, and RNA pull-down assays with assistance from J.L., Y.C., Y.Li., J.Y., Y.Liu., and L.W. L.W. conducted the HEK293T cell-based translation assays, while G.L. carried out the wheat germ extract and tobacco protoplast translation experiments. Cryo-EM grids were prepared by G.L. and Y.C. Cryo-EM data processing was performed by J.L. and G.L. Structural models were built and analyzed by J.L. and G.L. Data interpretation and manuscript writing were led by J.L. and G.L., with input from all authors. All authors reviewed and approved the final manuscript.

## Data availability

The coordinates and cryo-EM density maps for TLS^His^-60*S*, TLS^His^-80*S*, TLS^His^-80*S*-tRNA_i_^Met^, and CHX-TLS^His^-60*S* have been deposited under PDB and Electron Microscopy Data Bank accession codes 9VTG and EMD-65327; 9VTI and EMD-65329; 9VTH and EMD-65328; 9VTJ and EMD-65330, respectively.

