## Supplemental Information for "Viral tRNA-like structure hijacks host ribosomes for poly(A)-independent translation"

**Supplementary Methods**

***Preparation of RNAs***

To investigate TLS$^{His}$-mediated regulation of translation, firefly luciferase reporter constructs flanked by the TMV 5′ and 3′ UTRs were synthesized and cloned into the pcDNA3.1 vector backbone (GenScript). Various TLS$^{His}$ mutants was generated by GenScript, as listed in Supplementary Data 2. All *in vitro* transcribed mRNAs were synthesized by APExBio with a 5′ cap structure without modified nucleotides. Control mRNAs included uncapped transcripts, polyadenylated mRNAs (bearing a 100-nt poly(A) tail), and TLS$^{His}$-deleted constructs, in which the TMV 3′ UTR was removed.

To generate the isolated 204-nt TLS$^{His}$ element, a DNA fragment encoding the TMV 3′ UTR was amplified using a primer pair that incorporating a T7 promoter. The complete DNA sequence was:

5′-
GTAATACGACTCACTATA*GGTAGTCAAGATGCATAATAAATAACGGATTGTGTCC*
*GTAATCACACGTGGTGCGTACGATAACGCATAGTGTTTTTCCCTCCACTTAAATC*
*GAAGGGTTGTGTCTTGGATCGCGCGGGTCAAATGTATATGGTTCATATACATCCG*
*CAGGCACGTAATAAAGCGAGGGGTTCGAATCCCCCCGTTACCCCCGGTAGGGG*
*CCCA*-3′ (primers underlined, TLS$^{His}$ sequence in italics). The transcription reaction was assembled using standard HyperScribe T7 High Yield RNA Synthesis kit (ApexBio, K1074) and incubated at 37 °C for 3 h. The RNA was then purified via ion-exchange chromatography on a 1 mL Mono Q column, equilibrated in Q-buffer (50 mM Tris, pH 7.4, 80 mM KCl, 10 mM $MgCl_2$, 1 mM DTT) and eluted with a linear gradient to 1 M KCl. The RNA fragment was subsequently precipitated with 70% (v/v) ethanol and analyzed by urea-PAGE.

To label TLS$^{His}$ RNA, we employed a hybridization-ligation strategy. A 5′-labeled DNA oligonucleotide (5′-Cy5-GGGAGACCGGCAGATCT-3′ or biotin-labeled equivalent) was annealed to a bridging DNA strand (5′-ATTATGCATCTTGACTACCAGATCTGCCGGTCTCCC-3′), which was complementary to both the label and the 5′ end of the TLS$^{His}$ RNA. The TLS$^{His}$ RNA was first treated with T4 polynucleotide kinase (T4 PNK, NEB) to ensure a 5′ monophosphate group required for ligation. Then, the TLS$^{His}$ RNA, bridging oligo, and labeled oligo were mixed, heated to 80 °C for 2 min, and gradually cooled to 10 °C at a rate of 0.2 °C/s to promote proper annealing. A 40 μL ligation reaction was then assembled on ice, containing 4 μL 10× T3 DNA ligase buffer, 18 μM bridge DNA, 15 μM labeled DNA, 13 μM RNA, and 2 μL T3 DNA ligase (NEB). The reaction was incubated at 25 °C for 2 h. Following ligation, RNA products were purified by phenol–chloroform extraction and ethanol precipitation, and stored at −80 °C for downstream applications.

To generate a mini-genome RNA suitable for ribosome complex formation, a synthetic DNA construct was designed containing a 212-nucleotide segment derived from the 5′ end of the firefly luciferase ORF. This coding region was flanked by an 89-nt viral 5′ region (comprising a 68-nt 5′ UTR and the first seven codons of the viral sequence) and a 261-nt viral 3′ region encompassing the full-length $TLS^{His}$. The resulting DNA scaffold was cloned into a pET28a vector using *Xba*I and *Xho*I restriction sites. For in vitro transcription, 3 mM $m^7$GpppAmG (ApexBio, B8176) was added to the *in vitro* transcription reaction to generate capped RNAs bearing a Cap1 structure at the 5′ end. The full sequence of the synthetic mini-genome RNA is provided in table 1 of Supplementary Table.

***Preparation of tobacco ribosome***

Ribosome prepartion procudure discribed as previously[1], with minor modifications. Brifly, *Nicotiana benthamiana* plants were grown under physiological greenhouse conditions at 25 °C for 20 days. Leaf tissue was harvested and immediately flash-frozen in liquid nitrogen to preserve cellular integrity and arrest translation. Frozen tissue was ground to a fine powder using a pre-chilled mortar and pestle to mechanically lyse cells under RNase-free conditions. The powdered tissue was resuspended in plant extraction buffer containing 40 mM HEPES/ KOH pH 8.0, 100 mM KCl, 15 mM $MgCl_2$, 25 mM EGTA, 0.5 mg/ml heparin, 0.5 mM spermidine, 0.04 mM spermine, 10 mM DTT, 1% (w/v) Triton-100 and 2% (w/v) polyoxyethylene and supplemented with protease and RNase inhibitors.

The lystate was pelleted by ultracentrifugation over a 30% (w/w) sucrose cushion and centrifugation at 35,000 rpm for 20 h in a Beckman Type 45 Ti rotor, 4°C. The resulting ribosomal pellet was resuspended in dissociation buffer (20 mM HEPES, pH 8.0, 500 mM KCl, 2.5 mM $MgCl_2$, 1 mM DTT), followed by ultracentrifugation through a 10–40% linear sucrose gradient. Fractions were collected and analyzed by UV absorbance and mass spectrometry to confirm ribosomal subunit identity as listed in Supplementary Data 2.

***Polysome profiling***

Polysome profiling was performed following a modified protocol[2]. Approximately 2 g of flash-frozen *Nicotiana benthamiana* leaf powder was homogenized in 10 mL of polysome extraction buffer (200 mM Tris-HCl, pH 9.0, 200 mM KCl, 25 mM EGTA, 35 mM $MgCl_2$), supplemented with 0.2% (v/v) Brij-35, 0.2% Triton X-100, 0.2% Igepal CA-630, 0.2% polyoxyethylene 10 tridecyl ether, 5 mM DTT, and a protease inhibitor cocktail. The homogenate was sequentially filtered through double layers of Miracloth and clarified by centrifugation at 10,000 × g for 15 min at 4 °C. The supernatant was layered onto a 1.7 M sucrose cushion and subjected to ultracentrifugation at 170,000 × g for 3 h at 4 °C. The resulting ribosome pellet was resuspended in ribosome buffer (20 mM Tris-HCl, pH 8.0, 150 mM NaCl, 5 mM $MgCl_2$, 1 mM DTT).

For ribosomal profiling, 200 µL of ribosome preparation (corresponding to A260 ~ 2.0) was incubated with 1 µM final concentration of Cy5-labeled TMV 3′ UTR RNA (cy5-$TLS^{His}$) at room temperature for 30 minutes to allow complex formation. The mixture was gently layered onto a 10-60% (w/v) linear sucrose gradient prepared in ribosome buffer and subjected to ultracentrifugation at 46,000 rpm for 2 h at 4 °C using an SW60 rotor (Beckman Coulter). Gradient fractions were collected using a fractionator system (BioComp Instruments) equipped with inline UV monitoring to visualize ribosomal subunit distribution. RNA was extracted from each fraction by phenol-chloroform extraction followed by ethanol precipitation. Samples were resolved on 8% denaturing urea–PAGE gels, and Cy5 fluorescence signals were visualized by excitation at 600 nm to assess co-sedimentation of $TLS^{His}$ RNA with ribosomal complexes. For competition experiments, cycloheximide was added to a final concentration of 100 µg/mL prior to $TLS^{His}$ incubation. All other steps, including gradient separation and RNA analysis, were performed identically to the standard Cy5-$TLS^{His}$ profiling procedure.

***Ribosome pelleting***

Ribosome pelleting assays were conducted to evaluate the direct association between $TLS^{His}$ RNA and ribosomal subunits. Briefly, 20 µL binding reactions were assembled containing 1 µM purified human ribosomal subunit or 80*S*, 3 µM $TLS^{His}$ RNA, 50 mM Tris-acetate (pH 7.7), 50 mM potassium acetate, 3.8 mM magnesium acetate, and 10 mM DTT. The mixtures were incubated at room temperature for 15 min to allow binding and then carefully layered onto 200 µL of pre-chilled 32.5% (w/w) sucrose cushions. For competition assays with ribosome-targeting antibiotics, cycloheximide, puromycin, and homoharringtonine were pre-incubated with the ribosomes at a final concentration of 1 mM for 10 min at room temperature before adding $TLS^{His}$ RNA.

All samples were centrifuged at 38,000 rpm for 2 h at 4 °C using a Beckman Type 42.2 Ti rotor. After centrifugation, ribosomal pellets were gently washed and resuspended with ice-cold ribosome buffer. Associated RNAs were resolved by 8% denaturing urea-PAGE and visualized using Stains-All dye.

***RNA pull-down***

To isolate proteins that interact with the TMV $TLS^{His}$ element, we performed RNA pull-down assays using biotin-labeled $TLS^{His}$ RNA and streptavidin-coated magnetic beads (Dynabeads MyOne Streptavidin C1, Thermo Fisher). Freshly harvested BY-2 cells were lysed in five volumes of ice-cold extraction buffer (200 mM Tris-HCl, pH 9.0, 200 mM KCl, 25 mM EGTA, 35 mM $MgCl_2$, 0.2% Brij-35, 0.2% Triton X-100, 0.2% Igepal CA630, 0.2% polyoxyethylene 10 tridecyl ether, 5 mM DTT, and protease inhibitor cocktail). After clarification by centrifugation at 10,000 × g for 15 min at 4 °C, the supernatant was ready for RNA pull-down.

Prior to RNA immobilization, Dynabeads were washed three times with 2× binding and washing buffer (B&W: 20 mM Tris-HCl pH 7.5, 2 mM EDTA, 2 M NaCl), followed by incubation with biotin-labeled $TLS^{His}$ RNA (final concentration: 5 µg RNA per µL beads) in 1× B&W buffer at room temperature for 15 minutes with gentle rotation. After coupling, the RNA-coated beads were washed three times with 1× B&W buffer to remove unbound RNA. For each RNA pull-down assay, 50 µL of RNA-conjugated beads were incubated with 200 µL of freshly prepared cell lysate for 15 minutes at room temperature under gentle rotation. The beads were subsequently washed three times with resuspension buffer (20 mM Tris-HCl pH 8.0, 150 mM NaCl, 5 mM $MgCl_2$, 1 mM DTT). Bound proteins were eluted by heating the beads to 95 °C for 2 min in 1× B&W buffer, followed by centrifugation at 13,000 rpm for 2 minutes. The resulting supernatants were subjected to LC-MS/MS analysis on the institutional mass spectrometry platform to identify $TLS^{His}$-interacting proteins. Gene Ontology (GO) annotation of enriched proteins was performed using the DAVID bioinformatics tool[3].

***Tryptic digestion and LC-MS/MS analysis***

Protein samples were processed using a modified filter-aided sample preparation (FASP) protocol[4]. Briefly, samples were subjected to three rounds of buffer exchange with 8 M urea and 100 mM Tris-HCl (pH 8.0), followed by reduction with 10 mM dithiothreitol (DTT) at 37 °C for 30 min and alkylation with 30 mM iodoacetamide at 25 °C for 45 min in the dark. After three additional washes with digestion buffer (30 mM Tris-HCl, pH 8.0), tryptic digestion was performed overnight at 37 °C using trypsin at a 1:50 enzyme-to-protein ratio. The resulting peptides were collected by centrifugation, and filters were washed twice with 15% acetonitrile. All eluates were combined and dried under vacuum.

Peptide samples were resuspended and analyzed on an EASY-nLC 1200 system (Thermo Fisher Scientific) coupled to an Orbitrap Exploris 480 mass spectrometer. Peptides were separated on a home-packed C18 column (75 µm × 25 cm, ReproSil-Pur C18-AQ, 1.9 µm, Dr. Maisch GmbH) using a linear gradient of 5–44% mobile phase B (0.1% formic acid in 80% acetonitrile) over 38 min, followed by 44–70% B over 8 min, and 70–100% B over 2 min at a flow rate of 200 nL/min. High-field asymmetric waveform ion mobility spectrometry (FAIMS) was enabled with compensation voltages of −40 and −60 V. MS1 scans were acquired at 60,000 resolution, and data-dependent MS2 scans were performed in the Orbitrap at 15,000 resolution with a normalized collision energy (NCE) of 30%, using a 1.6 m/z isolation window and dynamic exclusion of 45 s.

Raw data were processed using Proteome Discoverer (v1.4, Thermo Fisher) with Mascot (v2.7.0, Matrix Science)[5] against the UniProt Nicotiana tabacum database (73,495 entries, downloaded April 29, 2025). Search parameters included a precursor mass tolerance of 10 ppm and a fragment mass tolerance of 0.05 Da, allowing up to two missed cleavages. Carbamidomethylation of cysteine was set as a fixed modification, while N-terminal acetylation and methionine oxidation were considered variable modifications.

***In vitro translation using Wheat Germ Extract***

In vitro translation assays were conducted using the wheat germ extract (WGE) translation system (Promega). The standard reaction mix was prepared in a total volume of 50 µL, which included 4 µL of amino acid mix (equally mixed amino acids), 5 µL of 3 M KOAc (pH 7.4), 25 µL of WGE. The reaction was initiated by adding 2.5 µL of RNA to a final concentration of 50 ng/µL. The reactions were performed in a thermoblock at 25 °C, and 10 µL of the reaction mix was taken at each time point and transferred to 40 µL of ice-cold 1× passive lysis buffer (Promega), followed by immediate freezing on dry ice. Luciferase activity was measured after thawing the samples at room temperature using a Varioskan lux system (thermo scientific) with firefly luciferase reagent (Promega). Data were analyzed with prism v9.0 using three replicate measurements for each time point.

#### *In vivo translation using HEK293T cell*

HEK293T cells were cultured in Dulbecco's Modified Eagle Medium (DMEM, Gibco) supplemented with 10% fetal bovine serum (FBS) at 37 °C in a humidified atmosphere with 5% $CO_2$. Cells were passaged every 72 hours using 0.2% trypsin-EDTA and reseeded onto 10 cm culture dishes.

For reporter assays, cells were seeded into 12-well plates at ~70% confluency. Transfection was performed using Lipofectamine 2000 (Invitrogen) according to the manufacturer's instructions. Briefly, 0.5 µg of *in vitro*–transcribed mRNA and 1.5 µL of Lipofectamine 2000 were each diluted in 100 µL Opti-MEM (Gibco), incubated separately for 5 min at room temperature, then combined and incubated for an additional 15 min. The transfection mixture was added dropwise to each well containing cells in fresh culture medium.

Cells were harvested at 2, 4, 8, 20, and 48 hr post-transfection. At each time point, cells were lysed in 100 µL of passive lysis buffer (Promega), vortexed for 30 s, and clarified by centrifugation at 12,000 × g for 2 min. For luminescence measurements, 20 µL of cell lysate was mixed with 100 µL of luciferase assay substrate and quantified using a Varioskan lux system.

#### *In vivo translation using tobacco protoplasts*

Suspension-cultured *Nicotiana tabacum* BY-2 cells were maintained in Murashige and Skoog (MS) medium supplemented with 30 g/L sucrose and 0.2 mL/L 2,4-dichlorophenoxyacetic acid (2,4-D, 1 mg/mL stock), adjusted to pH 5.8. Cultures were incubated in darkness at 25 °C with shaking at 130 rpm. For protoplast preparation, 3-day-old cells were harvested by centrifugation at 60 × g for 5 min and resuspended in enzyme digestion buffer containing 1.5% (w/v) Cellulase RS, 0.2% (w/v) macerase, 0.5 M mannitol, and 3.6 mM MES (pH 5.5). Digestion was performed in a 1 L flask at 28 °C with gentle shaking (100 rpm) for 4 hr. The resulting protoplasts were filtered through a 100 µm nylon mesh and washed twice with protoplast wash buffer (0.5 M mannitol, 3.6 mM MES, pH 5.5).

The pellet was resuspended in electroporation buffer (0.5 M mannitol, 3.6 mM MES, 60 mM KCl, pH 5.5), and the cell concentration was adjusted to $\sim 1 \times 10^6$ protoplasts/mL. For in vitro translation assays[6], 1 µg of in vitro–transcribed mRNA was added to 100 µL of freshly prepared protoplast suspension in electroporation buffer and gently mixed. Electroporation was performed using a single 400 V pulse in a Bio-Rad electroporation cuvette. Following electroporation, cells were immediately transferred to 12-well plates containing 800 µL protoplast culture medium (MS medium supplemented with 0.45 M mannitol and 50 µg/mL ampicillin) and incubated at 25 °C in the dark. To monitor translational kinetics, protoplasts were harvested at 2 h, 4 h, 8 h, 20 h, and 32 h post-transfection. For truncation series constructs, the culture medium was harvested 4 h after transfection. Luciferase reporter activity was measured following the same procedures described for HEK293T cell-based assays.

#### *cryo-EM sample preparation and data collection*

To investigate how $TLS^{His}$ engages the ribosome, we reconstituted 80*S*–$TLS^{His}$ complexes using purified Nicotiana benthamiana ribosomal subunits. A 20 µL in vitro assembly reaction was prepared containing 0.75 µM 40*S*, 0.75 µM 60*S*, 1.5 µM synthetic TMV mini-genome RNA, and 0.5 µL of yeast total tRNA (Sigma, R8508) in assembly buffer composed of 5 mM HEPES–NaOH (pH 7.5), 100 mM NaCl, 5 mM $MgCl_2$, and 5 mM β-mercaptoethanol. After 30 min incubation at room temperature, 2.5 µL of the reaction mixture was applied to glow-discharged graphene oxide-coated Quantifoil R2/1 300 mesh grids. Grids were blotted for 2-4 seconds (filter paper 595, TED Pella) under 100% humidity at 4 °C using a Vitrobot Mark IV (Thermo Fisher Scientific) and plunge-frozen in liquid ethane.

For cryo-EM sample preparation of the cycloheximide (CHX)-bound complex, a similar 20 µL reaction was assembled containing 0.75 µM 60*S*, 1.5 µM mini-genome RNA, and 1 mM CHX in the same assembly buffer, followed by grid preparation and vitrification using the same conditions as above.

Cryo-EM grids were loaded into a Thermo Scientific Titan Krios G4i transmission electron microscope equipped with a SelectrisX energy filter (10 eV slit width) and operated at 300 kV. Images were automatically acquired using EPU software with a Falcon 4i direct electron detector in EF-TEM mode at a nominal magnification of 130,000×, corresponding to a physical pixel size of 0.959 Å. The defocus range was set between −1.5 and −2.5 µm. Movies were recorded in EER format, with a total accumulated electron dose of ~50 $e^{-}/Å^{2}$. A detailed description of the data processing procedure is provided in the Supplementary Methods.

***cryoEM Data processing***

From 17,264 movies, a total of 3,455,822 particles were extracted following motion correction and CTF estimation[7]. After 2D classification, 1,642,799 particles were selected for ab initio reconstruction and heterogeneous refinement into five classes. The major classes corresponded to 60*S* and 80*S* ribosomal complexes, while a minor fraction represented junk particles. $TLS^{His}$ density was initially visible at the E site of both 60*S* and 80*S*, but global refinement was limited by local flexibility. To enrich for $TLS^{His}$-engaged particles, all 60*S* and 80*S* particles were merged and subjected to homogeneous refinement. A focused mask encompassing the ribosomal E and P sites—generated from a low-pass $TLS^{His}$ map and a molecular envelope of P-site-bound tRNA—was applied for 3D classification into 20 subclasses using signal subtraction. This strategy isolated particle subsets corresponding to $TLS^{His}$ bound to 60*S* alone, to free 80*S*, and to 80*S* containing initiator $tRNA_i^{Met}$, as well as classes showing tRNA occupancy in the P and/or E sites.

To reconstruct $TLS^{His}$-bound complexes at high resolution, particles from the enriched classes were re-extracted and subjected to homogeneous and non-uniform refinement[8]. For the $TLS^{His}$-60*S* complex, 53,930 particles yielded a map at 2.3 Å overall resolution, with local refinement around the $TLS^{His}$–L1 stalk interface improving resolution to 3.8 Å. The maps were merged using Phenix.combine_focused_maps to obtain composite models. For the $TLS^{His}$-80*S* complex, 422,599 particles provided a local resolution of 2.96 Å at the L1-$TLS^{His}$ region. The 80*S*-$TLS^{His}$-$tRNA_i^{Met}$ complex was reconstructed from 32,882 particles, yielding a local resolution of 4.88 Å.

Model building was initiated by docking the Nicotiana tabacum 80*S* ribosome (PDB: 8B2L)[1] into the 80*S*-$TLS^{His}$ map, followed by manual replacement of rRNA and ribosomal protein sequences with *N. benthamiana* homologs derived from transcriptomic data. The TLD region of $TLS^{His}$ was modeled manually into the 2.96 Å map. The UPD was generated using AlphaFold3[9] and fitted and refined into density as a rigid body. The complete models were iteratively refined in COOT[10] and Phenix real-space refinement[11] and combine focused maps. Final model validation and ADP refinement were performed prior to deposition. Statistics of map reconstruction and model quality are summarized in Supplementary Table 1. Figures and structural representations were prepared using UCSF Chimera[12] and ChimeraX[13].

### Supplementary Figures

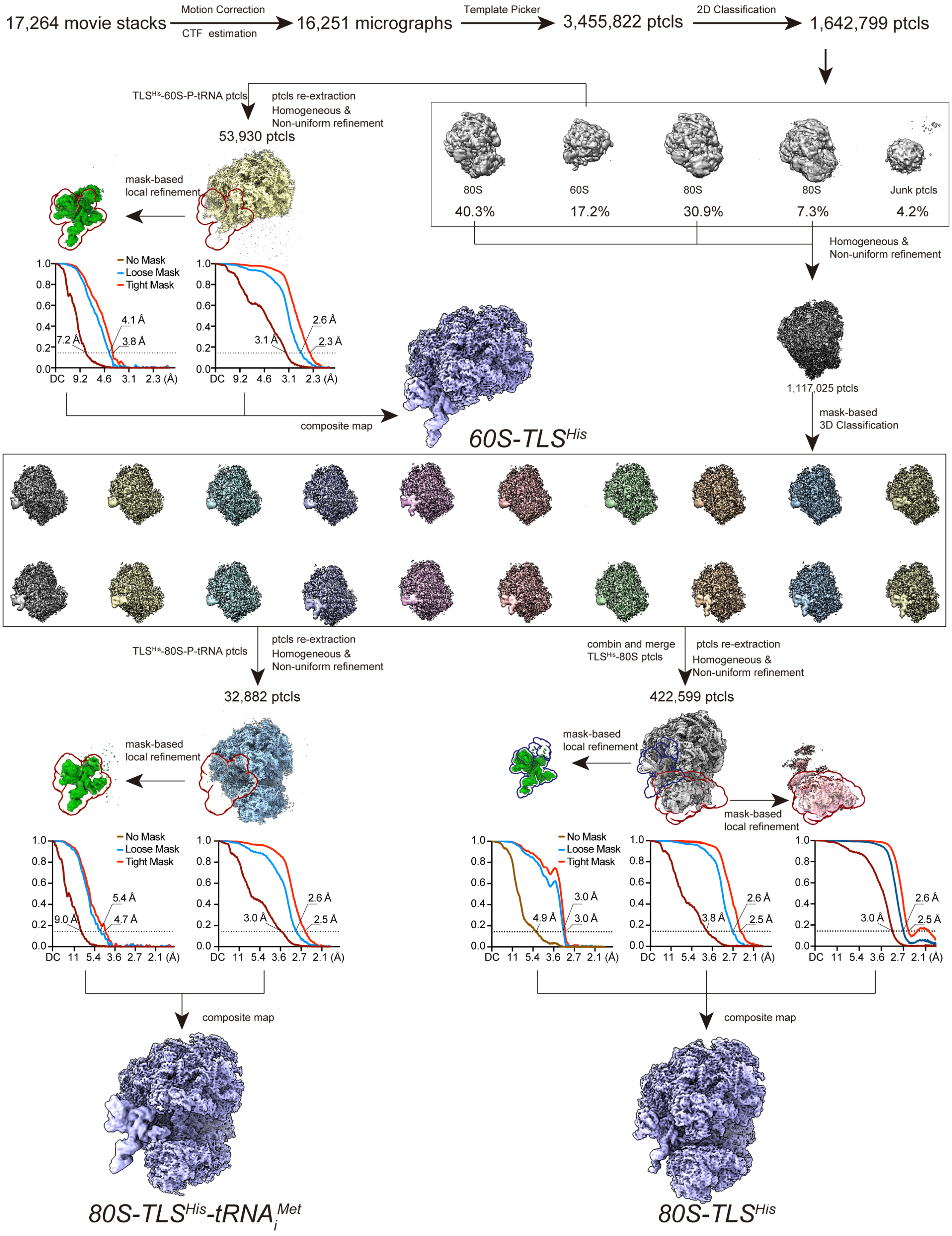


**Figure S1. Flowchart of the cryo-EM image processing and 3D reconstruction for TLS$^{His}$ complexes.**

Cryo-EM reconstruction of the TMV TLS–ribosome complex was performed in cryoSPARC. After *ab*-initio volume generation, iterative 3D classification progressively separated intact particles. A focused, mask-driven classification then enriched particles exhibiting well-defined P- and E-site densities. Subsequent masked refinement delivered a high-resolution map of the TLS$^{His}$-bound L1 stalk region. The final composite map was assembled in PHENIX by merging the best local reconstructions.

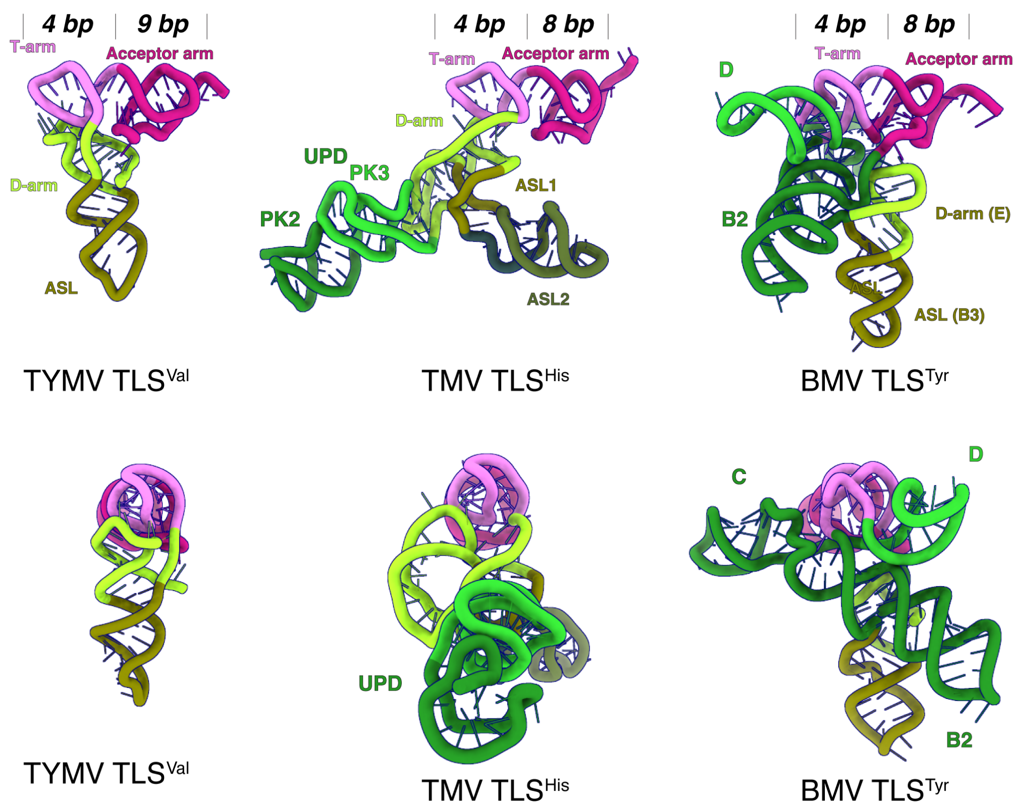


**Figure S2. Side-by-side comparison of three representative TLS structures.**

From left to right: $TLSv^{TYMV}$, $TLS^{HisTMV}$, and $TLSy^{BMV}$. All three TLSs fold into a canonical TLD plus an accessory module. In $TLS^{TYMV}$ and $TLS^{HisTMV}$, this accessory module sits upstream and governs translation; in $TLSy^{BMV}$, it is inserted within the TLD. The TLD itself comprises an acceptor arm and an anticodon-stem signature. A pronounced elbow, shaped by the T- and D-arms/loops, is evident in TLSv and $TLS^{His}$ but absent in TLSy. The number of base pairs in the T arm and acceptor arm is indicated above in italics. The color scheme matches that of Fig. 3.

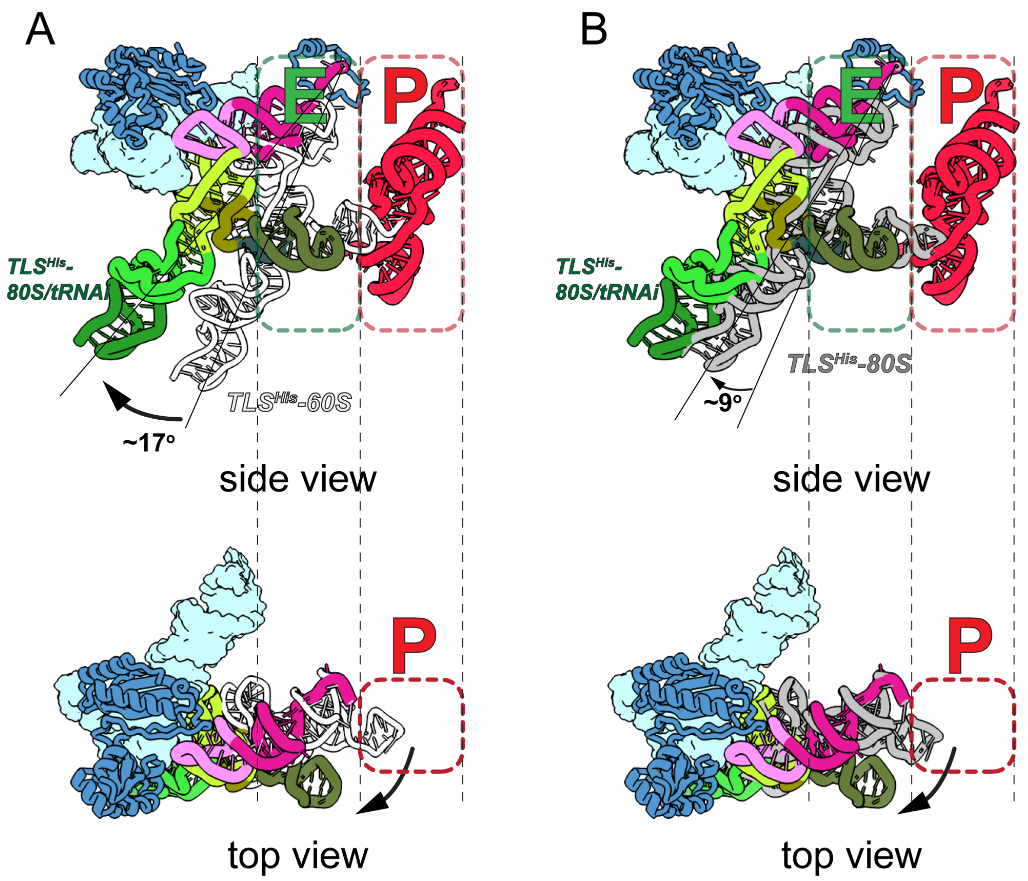


**Figure S3. Relative positioning of TLS$^{His}$ across different ribosome complexes, aligned by the 60*S* subunit.** (A) Structural comparison between the 60*S*-TLS$^{His}$ and 80*S*-TLS$^{His}$-tRNA$_i^{Met}$ complexes reveals a shift in TLS$^{His}$ orientation relative to the L1 stalk. (B) Comparison between TLS$^{His}$ bound in 80*S*-tRNA$_i^{Met}$ and free 80*S* complexes shows that TLS$^{His}$ undergoes an outward rotation of ~17° relative to the 60*S*-TLS$^{His}$ complex and ~9° relative to the 80*S*-TLS$^{His}$ complex, facilitating initiator tRNA accommodation at the P site. TLS$^{His}$ is shown in white for the 60*S*-TLS$^{His}$ complex and in gray for the 80*S*-TLS$^{His}$ complex. Color schemes follow those used in Fig. 3.

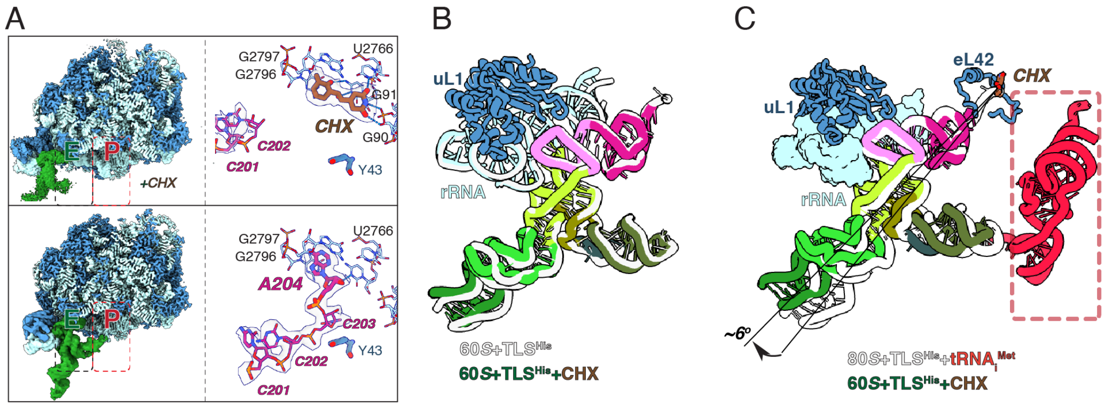


**Figure S4 TLS$^{His}$ bound to 60*S* in the presence of CHX.**
(A) Cryo-EM density comparison of 60*S*-TLSHis and CHX-60*S*-TLSHis; CHX occupies E-site overlapping TLS$^{His}$ CCA tail. (B), Superposition of CHX-bound and CHX-free 60*S*-TLS$^{His}$ using 25*S* rRNA (L1 stalk, residues 2454-2500) shows TLS$^{His}$ remains anchored. (C) Structural comparison of TLS$^{His}$ between 80*S*-tRNA$_i^{Met}$ and 60*S*-CHX-tRNA$_i^{Met}$ complexes. CHX-bound TLS$^{His}$ (color) shifts relative to CHX-free (white); rotation angles indicated.

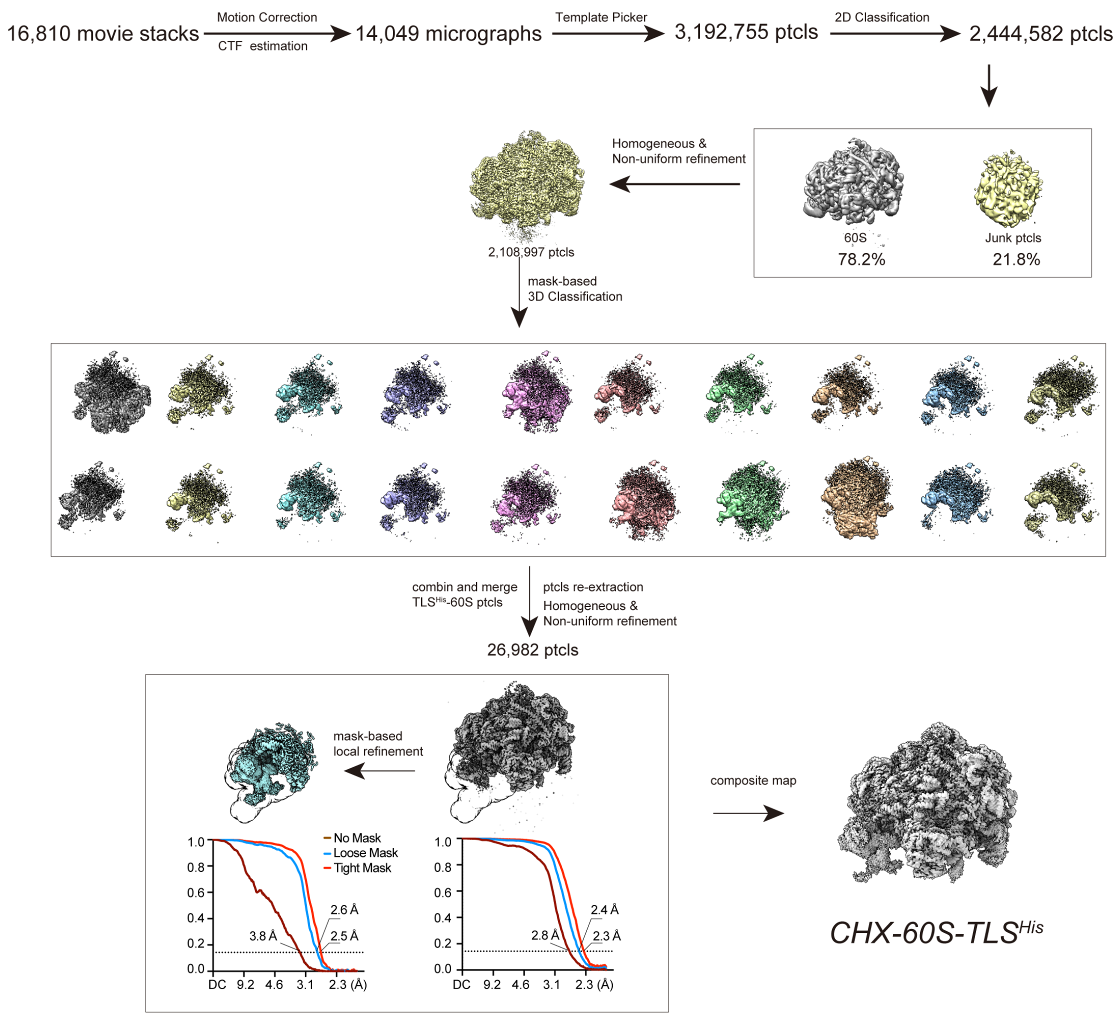


**Figure S5. Flowchart of cryo-EM data processing and 3D reconstruction for the TLS$^{His}$-60*S*-CHX complex.** The overall processing workflow mirrors that of Supplementary Fig. 1, with the key difference being an enlarged mask used during classification to encompass both the TLS$^{His}$ element and an extended portion of the 60*S* subunit.

**Table S1 Refinement and model statistics for TMV TLS$^{His}$ complexed with tobacco ribosome.**

| Complexes | TLS$^{His}$-60*S* | TLS$^{His}$-80*S* | TLS$^{His}$-80*S*-tRNA$_i^{Met}$ | TLS$^{His}$-60*S*-CHX |
|---|---|---|---|---|
| PDB and EMDB | 9VTG/EMD-65327 | 9VTI/EMD-65329 | 9VTH/EMD-65328 | 9VTJ/EMD-65330 |
| **Data collection**[a] | | | | |
| Magnification | 130,000x | 130,000x | 130,000x | 130,000x |
| Voltage (kV) | 300 | 300 | 300 | 300 |
| Total dose ($e^-/Å^2$) | 50 | 50 | 50 | 50 |
| Defocus range (μm) | -1.5 to -2.5 | -1.5 to -2.5 | -1.5 to -2.5 | -1.5 to -2.5 |
| Pixel size (Å) | 0.959 | 0.959 | 0.959 | 0.959 |
| Automation software | EPU | EPU | EPU | EPU |
| Symmetry imposed | C1 | C1 | C1 | C1 |
| **Reconstruction** | | | | |
| Number of used micrographs | 16,251 | 16,251 | 16,251 | 14,049 |
| Total of extracted particles | 1,642,799 | 1,642,799 | 1,642,799 | 3,192,755 |
| Total of refined particles (no.) | 53,930 | 422,599 | 32,882 | 26,982 |
| Resolution Masked FSC 0.143 | 3.8 Å | 3.0 Å | 4.7 Å | 2.5 Å |
| **Model Refinement** | | | | |
| Non-hydrogen atoms | 129,544 | 199,397 | 201,638 | 129,498 |
| Bond lengths (Å) | 0.007 | 0.007 | 0.007 | 0.008 |
| Bond angles (°) | 1.009 | 0.973 | 0.961 | 1.009 |
| **Validation** | | | | |
| MolProbity score | 1.59 | 1.74 | 1.71 | 1.62 |
| All-atom clashscore | 3.63 | 4.43 | 4.67 | 3.73 |
| Rotamers outliers (%) | 2.71 | 3.24 | 3.10 | 2.91 |
| Cβ outliers (%) | 0.00 | 0.01 | 0.00 | 0.00 |
| CaBLAM outliers (%) | 1.16 | 1.26 | 1.20 | 1.16 |
| B-factors (min/max/mean) | | | | |
| Protein | 1.00/231.19/28.74 | 1.00/299.04/71.46 | 1.00/348.97/333.45 | 5.56/127.05/22.07 |
| Nucleotide | 1.00/629.17/42.02 | 1.00/228.37/56.54 | 1.00/853.42/40.29 | 1.83/282.35/46.92 |
| Overall correlation coefficients | | | | |
| CC (mask) | 0.78 | 0.82 | 0.70 | 0.74 |
| CC (peaks) | 0.69 | 0.76 | 0.60 | 0.64 |
| CC (volume) | 0.78 | 0.80 | 0.69 | 0.74 |
| Ramachandran plot statistics | | | | |
| Favored (%) | 97.46 | 97.33 | 97.53 | 97.44 |
| Allowed (%) | 2.51 | 2.64 | 2.46 | 2.54 |
| Disallowed (%) | 0.03 | 0.04 | 0.01 | 0.02 |

[a] one data set from a single grid was used for data collection and processing for 60*S*-TLS$^{His}$, 80*S*-TLS$^{His}$, and 80*S*-tRNA$_i^{Met}$-TLS$^{His}$; one data set for 60*S*-TLS$^{His}$-CHX.
